# CHOmpact: a reduced metabolic model of Chinese hamster ovary cells with enhanced interpretability

**DOI:** 10.1101/2021.07.19.452953

**Authors:** Ioscani Jiménez del Val, Sarantos Kyriakopoulos, Simone Albrecht, Henning Stockmann, Pauline M Rudd, Karen M Polizzi, Cleo Kontoravdi

## Abstract

Metabolic modelling has emerged as a key tool for the characterisation of biopharmaceutical cell culture processes. Metabolic models have also been instrumental in identifying genetic engineering targets and developing feeding strategies that optimise the growth and productivity of Chinese hamster ovary (CHO) cells. Despite their success, metabolic models of CHO cells still present considerable challenges. Genome scale metabolic models (GeMs) of CHO cells are very large (>6000 reactions) and are, therefore, difficult to constrain to yield physiologically consistent flux distributions. The large scale of GeMs also makes interpretation of their outputs difficult. To address these challenges, we have developed CHOmpact, a reduced metabolic network that encompasses 101 metabolites linked through 144 reactions. Our compact reaction network allows us to deploy multi-objective optimisation and ensure that the computed flux distributions are physiologically consistent. Furthermore, our CHOmpact model delivers enhanced interpretability of simulation results and has allowed us to identify the mechanisms governing shifts in the anaplerotic consumption of asparagine and glutamate as well as an important mechanism of ammonia detoxification within mitochondria. CHOmpact, thus, addresses key challenges of large-scale metabolic models and, with further development, will serve as a platform to develop dynamic metabolic models for the control and optimisation of biopharmaceutical cell culture processes.

## 1. Introduction

Production of recombinant proteins is known to compete with biomass synthesis for externally provided nutrients. This is particularly true for mammalian cell lines, such as Chinese hamster ovary (CHO) cells, which are the dominant host for industrial production of therapeutic proteins (O’Flaherty et al., 2020). Metabolic modelling has become an essential tool for understanding resource allocation and, coupled with advances in genome editing, designing rational cell engineering strategies. Publication of the CHO-K1 genome, and the omics analyses this enabled, laid the foundation for systems-level understanding of this host. This knowledge has been reconstructed mathematically in a community-curated genome-scale metabolic model (GeM) of the CHO cell termed iCHO1766 (Hefzi et al., 2016). Crucially, the GeM organised knowledge of all biochemical conversions, transport and exchange reactions to create a large, interlinked network of metabolites and their associated reactions.

The inclusion of gene-protein reaction associations provided a direct link between genes and metabolic reactions. Since then, significant expansions and improvements to iCHO1766 have been achieved, such as gap-filling studies that also removed dead-end reactions (Fouladiha et al., 2021), and the integration of a core protein secretory pathway, iCHO2048, enabling the computation of energetic costs and machinery demands of each secreted protein (Gutierrez et al., 2020). Interestingly, iCHO2048 was subsequently used to direct host cell protein knockout studies which resulted in increased recombinant protein productivity and a cleaner feedstock for downstream processing steps (Kol et al., 2020), highlighting the power these models hold for identifying diverse cellular engineering strategies.

The solution of GeMs, and any undetermined metabolic model, relies on constraint-based methods, such as flux balance analysis (FBA), to predict steady-state intracellular flux distributions (Orth et al., 2010). Although FBA offers the advantage of not requiring detailed knowledge of enzymatic kinetic parameters, it does not return a unique set of intracellular flux values. In addition, the larger the metabolic network considered, the more difficult it becomes to interpret such predictions (Gardner and Boyle, 2017). GeMs therefore require large datasets, preferably across different omics levels (e.g., metabolomic, transcriptomic) to increase confidence in results. This is also true for curating GeMs for specific cell lines or systems, raising the need for extensive experimentation that goes beyond typical analytical measurements conducted in an industrial setting.

Several algorithms have been developed to improve the predictive performance of GeMs, for example by constraining the amount of carbon able to flowthrough reaction fluxes, based on the maximum amount carbon uptake by the cell (ccFBA) (Lularevic et al., 2019), taking into account the selective pressure that exists within cell cultures for fast-growing cell lines with a low enzyme usage (Lewis et al., 2010), or introducing enzyme capacity constraints (Yeo et al., 2020). Despite these advances, both the accuracy and interpretability of intracellular flux predictions remain challenging. An additional limitation is the computational difficulty in creating dynamic versions of GeMs that would reflect the nature of cell culture processes, although recent efforts coupling a CHO GeM with statistical models have yielded promising results in predicting the time evolution of extracellular amino acid concentrations (Martínez et al., 2015).

In this work, we introduce a reduced-scale metabolic model, CHOmpact, where the reaction network is based on the work by Carinhas et al. (2013) and has been augmented with a detailed description the aspartate-malate (Asp-Mal) shuttle, the urea cycle, de novo serine synthesis from glycolytic intermediates, and nucleotide sugar donor biosynthesis. The resulting network comprises 101 metabolites and 144 reactions, which due to its compact nature, significantly enhances the interpretability of simulation results. The reduced scale of the network and associated FBA problem also allow for more complex, non-linear formulations of the objective function to be incorporated compared to biomass maximisation that is often employed in FBA of GeMs. The multi-objective optimisation framework used to solve CHOmpact allows us to solve across all phases of cell culture and provides insight into the dynamics of cellular metabolism. We envisage that the advantages presented by CHOmpact will enable the development of dynamic flux balance models that can serve as digital twins for the control and optimisation of biopharmaceutical cell culture processes.

## 2. Materials & Methods

### 2.1. Experimental

#### 2.1.1. Cell culture

The GS46 GS-CHO cell line producing a humanized anti-Tumour-Associated Glycoprotein (TAG-72) IgG4κ mAb (cB72.3), a kind gift by Lonza Biologics (Slough, UK), was cultured with three different amino acid feeds: Feed C, Feed U and Feed U40 (Kyriakopoulos and Kontoravdi, 2014). Briefly, triplicate cultures for each feeding regime were performed in orbitally shaken (140 rpm) 250mL vented conical flasks (Corning, Amsterdam, Netherlands) with a 50mL working volume. The cultures were performed in a humidified incubator with CO_2_ controlled at 8% and temperature set at 36.5°C. The basal culture medium for all cultures was CD CHO (Life Technologies, Paisley, UK) supplemented with 25μM methionine sulfoximine (Sigma-Aldrich, Dorset, UK). All feeding regimes consisted in adding 10% v/v every 48 hours of culture starting on day 2. The Feed C regime used commercial CD EfficientFeed™ C AGT™ (Invitrogen, UK), whereas the U and U40 feeds aimed to provide growth-limiting nutrients (glucose and amino acids) beyond the amounts available in Feed C. The glucose and amino acid concentrations present in the different feeds is detailed in previous work by Kyriakopoulos and Kontoravdi (2014).

#### 2.1.2. Analytical methods

Viable and dead cell density was determined using the trypan blue dye exclusion method and light microscopy. mAb titre was determined using the BLItz® system (Pall ForteBio, Portsmouth, UK). Time profiles for glucose, lactate, and ammonia were generated using the Bioprofile 400 analyser (NOVA Biomedical, Waltham, MA). Residual amino acid profiles were generated using the PicoTag method (Waters, Hertfordshire, UK) on an Alliance HPLC instrument (Waters, Hertfordshire, UK). Extracellular pyruvate concentrations were determined with an enzyme assay kit (Abcam, Cambridge, UK). mAb Fc glycoprofiling was performed with an automated sample preparation workflow (Stockmann et al., 2013). Briefly, the mAb samples were affinity-purified from the cell culture supernatant with a 96-well Protein G IgG purification plate (Thermo Fisher Scientific, Dublin, Ireland). Glycans were released from the mAb through PNGase (Prozyme, Hayward, California) digestion and labelled with 2-amino benzamide (Ludger, Oxford, UK). Labelled glycans were separated using ultra-performance hydrophilic interaction chromatography (UPLC-HILIC) and quantified through fluorescence detection (Stockmann et al., 2013). Glycans were initially assigned by comparing their Glucose Unit retention times with those available in the NIBRT GlycoBase 3.2 structural N-glycan library (Campbell et al., 2008). Glycan assignment was confirmed through weak anion exchange chromatography and quadrupole time-of-flight mass spectrometry on exoglycosidase-digested samples (Albrecht et al., 2014).

#### 2.1.3. Dry cell weight measurement

Cells were cultured under the same conditions as described for Feed C, above. Duplicate cultures were harvested at day 4 and day 10 for dry cell weight (DCW) measurements for cells undergoing mid-exponential and stationary growth, respectively. Prior to harvest, viable cell density was determined using the trypan blue dye exclusion method. Immediately after cell counting, 40mL of the cultures was harvested and centrifuged at 1000g for one minute in pre-weighed 50mL falcon tubes. The supernatant was then discarded, and the cell pellets were washed once with 40mL 0.9% w/v NaCl (Sigma-Aldrich, Dorset, UK) solution and centrifuged at 1000g for one minute. The wash was discarded, and the cell pellet was left to dry in a non-humidified incubator at 37°C until no changes in weight were observed. The tubes were weighed within 1mg accuracy (ACCULAB, Sartorius, Surrey, UK).

#### 2.1.4. Data processing and analysis

The cell-specific rates for nutrient consumption and metabolite/product secretion, *q*_*i*_ (*t*_*n*_), were calculated by performing linear regressions to obtain the slopes in Eq. 1 (Sauer et al., 2000), where *N*_*i,cons*_(*t*_*n*_) is the consumed/produced amount of component *i* (in units of *nmol*_*i*_ or *mg*_*mAb*_) up to time *t*_*n*_ and *IVC*(*t*_*n*_) is the integral of viable cells up to time *t*_*n*_ in units of 10^6^ cells h.

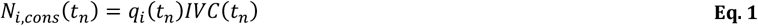

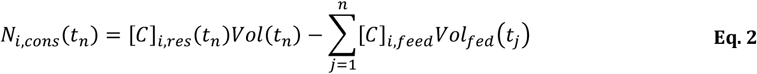

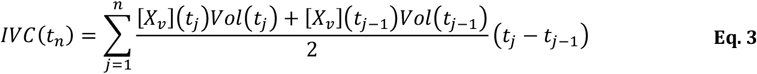

*N*_*i,cons*_(*t*_*n*_) and *IVC*(*t*_*n*_) were computed using Eq. 2 and Eq. 3, respectively. In Eq. 2, [*C*]_*i,res*_(*t*_*n*_) is the residual concentration of component *i* at time *t*_*n*_, [*C*]_*i,feed*_ is the concentration of component *i* in the feed and *Vol*_*fod*_ (*t*_*j*_)is the feed volume added at time *j*. In Eq. 3, [*X*_*v*_](*t*_*j*_)is cell density at time *t*_*j*_ and *Vol*(*t*_*j*_) is liquid volume in the culture flask at time *t*_*j*_ .

The linear regressions for the determination of cell specific uptake rates were computed using the LINEST function in Microsoft Excel. Confidence intervals for the obtained *q*_*i*_ (*t*_*n*_) values were computed with the least square residuals and a t-value for *p* = 0.05. Data analysis identified six distinct intervals with constant uptake/secretion rates: early exponential, mid-exponential, late exponential, early stationary, and stationary. Raw cell culture data is presented in Supplementary Figure 1, and the processed data, along with the identified intervals are presented in Supplementary Figure 2.

The amount of mAb glycoform secreted across different intervals was calculated using Eq. 4, where *f*_*k*_ represents the fraction of mAb glycoform *k* secreted during the time interval (*t*_*n*_ − *t*_*n*−1_), [*mAb*]_*i*_ (*t*_*n*_) is the concentration of mAb glycoform *k* present at time *t*_*n*_ and [*mAb*](*t*_*n*_) is the total mAb titre at time *t*_*n*_ (del Val et al., 2016a; Fan et al., 2015).

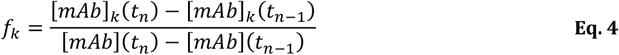

### 2.2. Flux balance model development

Our flux balance model (Figure 1) is based on previous work for GS-CHO cells (Carinhas et al., 2013), which has been expanded to include the aspartate-malate (Asp-Mal) shuttle (Mulukutla et al., 2012; Nolan and Lee, 2011), the urea cycle (Zamorano et al., 2010), *de novo* serine synthesis from glycolytic intermediates, and ATP synthesis via oxidative phosphorylation. The pathway for nucleotide sugar donor biosynthesis (Kremkow and Lee, 2018) has also been included.

**Figure 1.**
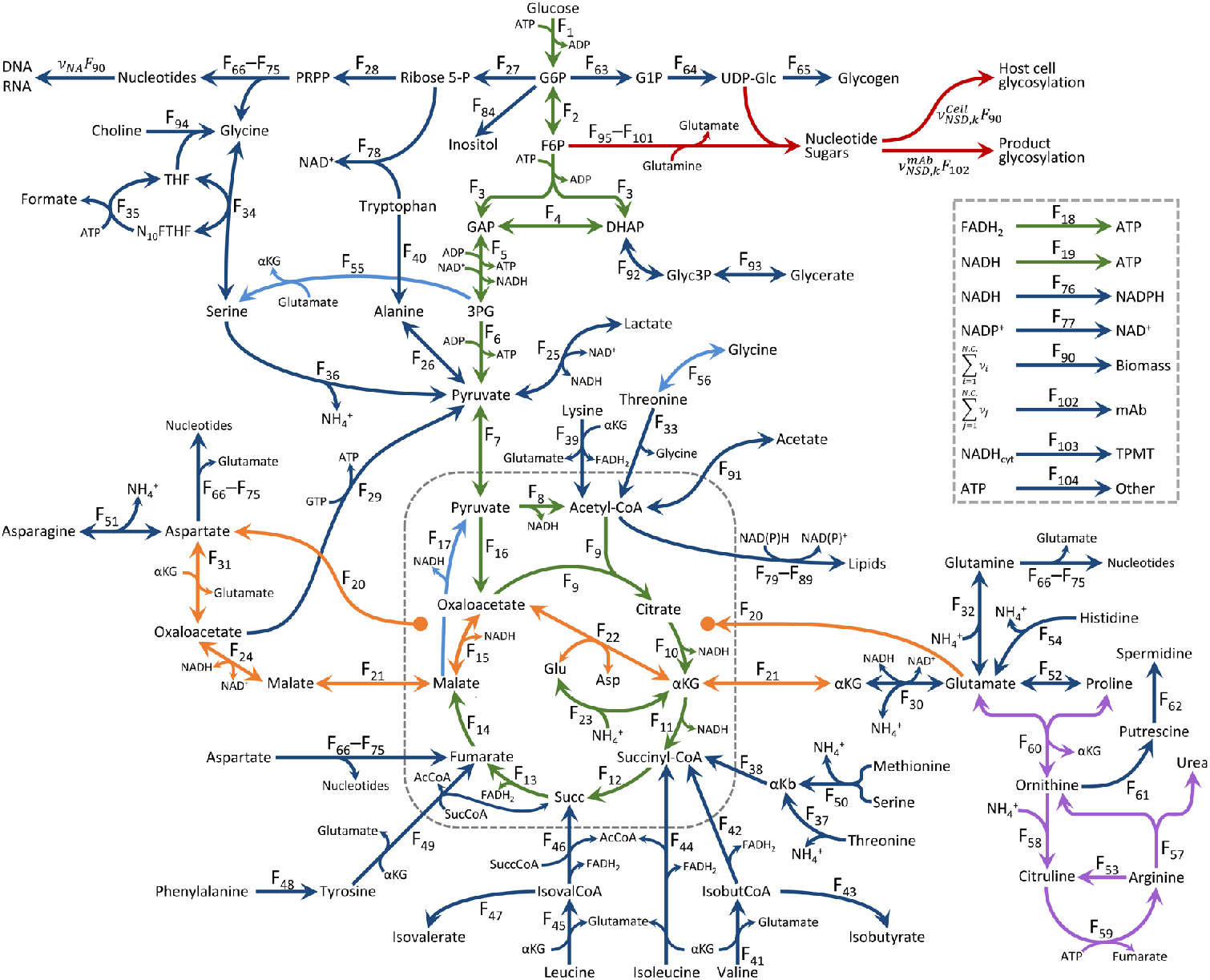
FBA reaction network. The present FBA model considers 101 species linked through 144 fluxes. Different colours indicate particular metabolic pathways: glycolysis, TCA and oxidative phosphorylation (Green), nucleotide sugar donor metabolism (Red), aspartate-malate shuttle (Orange), urea cycle (Purple), amino acid and nucleotide metabolism (Dark Blue) and cycle fluxes that must be constrained and/or estimated during optimisation (Light Blue).

Manual curation of our FBA model was performed using the KEGG database (Kanehisa et al., 2017; Kanehisa et al., 2019) and the reference CHO-K1 and *Cricetulus griseus* genome annotations (Kremkow et al., 2015; Lewis et al., 2013; Rupp et al., 2018). The resulting model comprises 101 metabolites linked through 299 reactions (Supplementary Table 1). Supplementary Table 1 also provides links for all enzymatic reactions to KEGG (Kanehisa et al., 2017; Kanehisa et al., 2019) as well as to reference sequences in the NCBI database (O’Leary et al., 2016).

Sequential reactions throughout the metabolic pathways were combined into single reaction fluxes to reduce degrees of freedom within the model (Nolan and Lee, 2011). Overall, the resulting metabolic model considers material balances for 101 species, one additional equation that defines the consumption of ATP towards active amino acid transport and 144 fluxes (Figure 1 and Supplementary File, Section 2), yielding 42 degrees of freedom. The full stoichiometric matrix underlying our model is presented in Supplementary Table 2.

#### 2.2.1. Stoichiometric equations for biomass and product

Calculations for the biomass stoichiometric coefficients are presented in Supplementary Table 3. Based on experimental measurements, the biomass stoichiometric equation considers a dry cell weight of 219 pg/cell for exponentially growing cells and 311 pg/cell for cells in stationary phase. The mass composition of GS-CHO cells is based on Sheikh et al. (2005) and Hefzi et al. (2016) and assumes 74.2% protein, 11.1% lipids, 5.0% RNA, 1.4% DNA, 0.4% glycogen, 0.2% N-glycans, 0.3% O-glycans, 2.9% other intracellular components (e.g. MTHF, NAD(P)H, AcCoA) and 4.5% non-balanced components (ash).

The individual amino acid content of protein was computed from CHO cell proteomic data (Baycin-Hizal et al., 2012), as reported previously (del Val et al., 2016b). The glycan content of biomass, which has been included by using the biomass NSD stoichiometric coefficients for cellular protein N- and O-linked glycosylation as well as glycolipid glycosylation (del Val et al., 2016b).

The stoichiometric equation for the cB72.3 product, a humanised IgG4κ mAb, was computed based on the amino acid sequences for the human IgG4 Fc (Heilig et al., 2003), the constant fragment of a human kappa light chain (Brady et al., 1991; Xiang et al., 1999), as well as the variable heavy and light chain fragments for the cB72.3 mAb (Xiang et al., 1999). The sequences and calculations for the mAb amino acid stoichiometric coefficients are presented in Supplementary Table 4.

mAb glycoprofiling at three culture timepoints (192h, 240h, and 288h) allowed us to calculate stoichiometric coefficients for NSD consumption towards mAb glycosylation across three culture intervals: 0 to 192 hours, 192 to 240 hours, and 240 to 288 hours. These calculations were made with Eq. 4, and the obtained stoichiometric values are presented in Supplementary Table 5.

#### 2.2.2. FBA solution: multi-objective optimisation

As with most FBA models, no intracellular accumulation of species has been assumed in the material balances generated from our stoichiometric matrix, leading to a problem of the form:

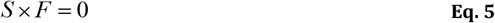

Where *S* is the stoichiometric matrix defined in Supplementary Table 2 and *F* is the vector of unknown fluxes. Because the model contains more unknown fluxes (144) than equations (102), it must be solved using constraint-based optimisation strategies that have been outlined elsewhere (Banga, 2008).

Two constraint-based optimisation strategies have been used to solve our FBA. The first is typical in that it maximises the rate of biomass synthesis while maintaining the transport flux for all nutrients, metabolites, and product set to their experimentally determined values. Reaction reversibility constraints, based on enzyme data available in the KEGG (Kanehisa et al., 2017; Kanehisa et al., 2019) and BRENDA (Jeske et al., 2019) databases, were included and are indicated in Supplementary Tables 1 and 2.

The multi-objective optimisation strategy proposed herein simultaneously maximises the fluxes where ATP is synthesised while minimising the sum of squared intracellular fluxes. This objective function represents maximum energetic efficiency by the cells (Schuetz et al., 2007) and was used to ensure consistent directionality of central carbon metabolism fluxes. Alongside maximising the energetic efficiency of the cells, the squared difference between measured and computed fluxes was minimised to ensure consistency between our flux model results and our experimental measurements. In order to avoid flux *F*_15_ being bypassed by *F*_17_, the *F*_17_/*F*_14_ ratio was constrained to values within the [0, 1] interval and minimised. Finally, the sum of non-measured by-product secretion fluxes was also minimised as part of this multi-objective optimisation strategy. As with the traditional optimisation strategy, flux reversibility constraints were included within our multi-objective optimisation strategy.

An additional constraint was included for the maximum allowable fluxes along the aspartate-malate (Asp-Mal) shuttle. Flux through the Asp-Mal shuttle was constrained by limiting the maximum amount of glutamate transported into the mitochondrial lumen (*F*_20_) to the flux of glutamate internalised by the cells or produced through reactions independent of the Asp-Mal shuttle. *F*_20_ was constrained because this antiport transport flux is the rate-limiting step of the Asp-Mal shuttle (LaNoue et al., 1974; LaNoue and Tischler, 1974).

Importantly, a constraint for Asp-Mal fluxes is required because the sum of all reactions in the shuttle result, exclusively, in net transport of cytosolic NADH into the mitochondrial lumen (see Supplementary File). Thus, the fluxes through the reactions underlying the Asp-Mal shuttle can take any value as long as the balances for cytosolic and mitochondrial NADH are met (i.e., all other fluxes cancel each other out). This is likely to be the cause of what may be considered high (and possibly inconsistent) Asp-Mal shuttle fluxes with values that are comparable with glucose uptake fluxes (Mulukutla et al., 2012; Nolan and Lee, 2011). Avoiding these potential inconsistencies led us to include this Asp-Mal constraint within our FBA solution strategy.

Our multi-objective optimisation strategy also constrains *F*_16_ to be 1% of the glucose uptake flux (*F*_105_), as previously determined for CHO cells undergoing both exponential and stationary growth (Ahn and Antoniewicz, 2013). Both optimisation strategies are outlined in Table 1, where *F*_*j*_ are all intracellular fluxes, *ATP*_*synth* ._ is the squared sum of ATP synthesis reaction fluxes, *SSE* is the sum of square errors between the measured and computed transport fluxes, *F*_*K* ,*BP*_ are the fluxes of by-product synthesis reactions, *LB*_*i*_ and *UB*_*i*_ are lower and upper bounds for flux values (a reaction is irreversible when *LB*_*i*_ = 0), *C*_*Asp* /*Mal*_ is the Asp-Mal shuttle constraint, ε represents a small threshold value, 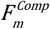 and 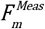 are the computed and measured transport fluxes, respectively, *F*_*g*_ are the fluxes of reactions where Glu is produced or consumed (excluding Asp-Mal shuttle reactions), and *v*_*g*_ is the stoichiometric coefficient for these Glu synthesis reactions.

**Table 1.** Optimisation strategies for flux balance model solution

| BM maximisation | Multi-objective optimisation |
| --- | --- |
| $MAX(F_{142})$<br>Subject to:<br>$\begin{cases} LB_i \leq F_i \leq UB_i \\ \left[ 100 \left( \frac{F_{143}}{q_p} - 1 \right) \right]^2 \leq \varepsilon_p \end{cases}$ | $MIN \left( \frac{\sum_{j=1}^{104} F_j^2}{ATP_{synth.}} + SSE + \sum_{k=1}^{14} F_{k,BP}^2 + \frac{F_{17}}{F_{14}} \right)$<br>Subject to:<br>$\begin{cases} LB_i \leq F_i \leq UB_i \\ 0 \leq \frac{F_{17}}{F_{14}} \leq 1 \\ C_{Asp/Mal} \leq \varepsilon_{Asp/Mal} \\ \left[ 100 \left( \frac{F_{16}}{F_{105}} - 0.01 \right) \right]^2 \leq \varepsilon_{16/105} \end{cases}$<br>Where:<br>$ATP_{synth.} = (F_5)^2 + (F_6)^2 + (1.5F_{18})^2 + (2.5F_{19})^2$ $SSE = \sum_{k=1}^{38} \left[ 100 \left( \frac{F_m^{Comp.}}{F_m^{Meas.}} - 1 \right) \right]^2$ $C_{Asp/Mal} = \left( \sum_g^{18} v_g F_g - F_{20} \right)^2$ |

The biomass maximisation strategy was used to compare performance of our reduced FBA model with the iCHO1766 GeM (Hefzi et al., 2016) while the energetic efficiency maximisation strategy was used for all other simulations presented herein. All optimisations were performed using the nonlinear programming sequential quadratic programming (NLPSQP) solver built into gPROMS ModelBuilder v6.0.2 (Process Systems Enterprise) on a standard desktop workstation (AMD Ryzen 2700x @ 4GHz and 16GB RAM).

## 3. Results and discussion

### 3.1. The reduced reaction network performs comparably with the iCHO1766 GeM

The maximum specific growth rates (*μ*_*g,max*_) of multiple CHO cell lines cultured under different conditions were calculated through the biomass synthesis rate maximisation strategy. The required inputs for this solution strategy, namely the nutrient, metabolite, and product transport fluxes, were obtained from previously published data (Carinhas et al., 2013; Martínez et al., 2015; Selvarasu et al., 2012), as summarized in the supplementary information of Hefzi et al. (2016).

In order to assess whether our assumed biomass composition and reduced reaction network performs comparably with a large-scale model for CHO cell metabolism, Figure 2 compares our predicted *μ*_*g,max*_ values with those obtained using the iCHO1766 GeM (Hefzi et al., 2016) and the corresponding experimental data.

**Figure 2.**
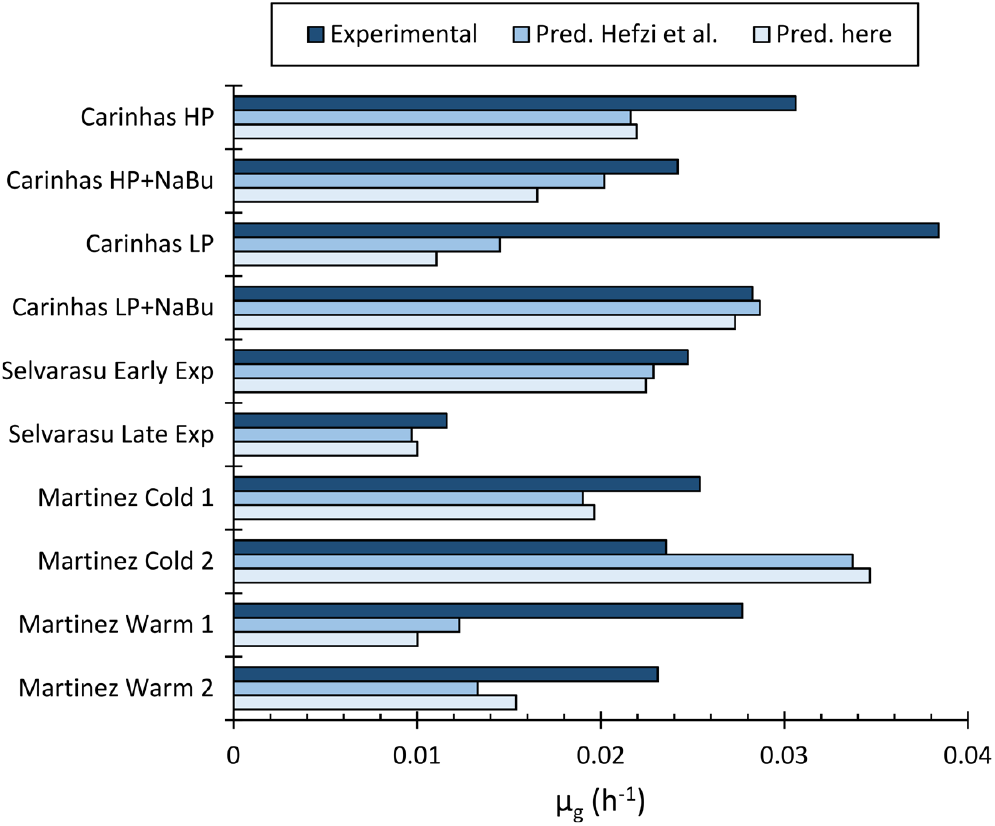
Comparison of experimentally determined and predicted maximum specific growth rates (*μ*_*g,max*_) for different CHO cell lines and culture conditions. The dark blue bars present the experimentally determined *μ*_*g,max*_ values reported by Carinhas et al. (2013), Martínez et al. (2015), and Selvarasu et al. (2012). The medium blue bars present *μ*_*g,max*_ predictions reported by Hefzi et al. (2016) and the light blue bars show the *μ*_*g,max*_ values predicted with CHOmpact.

Figure 2 shows that our model accurately predicts (within 15% deviation) the experimental data for the Low Producing CHO cell line cultured with sodium butyrate (LP+NaBu) from Carinhas et al. (2013) as well as the early and late exponential growth phases reported by Selvarasu et al. (2012). Our model underpredicts the growth rate reported for the High Producing CHO cell line cultured in absence and presence of NaBu (HP and HP+NaBu, respectively) (Carinhas et al., 2013) as well as CHO cells cultured under normal and mild hypothermic conditions (Warm 1, Warm 2, and Cold 1, respectively) (Martínez et al., 2015). Our model overpredicts the biomass synthesis rate of the second hypothermic dataset (Cold 2) reported by Martínez et al. (2015).

*μ*_*g,mAb*_ optimisations were performed using the biomass composition assumed by Hefzi et al. (2016) in order to discern whether the deviation in predictive capabilities of our model were caused by our assumed biomass composition. These optimisations yield overall average deviations from the experimental data that are indistinguishable from those obtained with our assumed biomass composition (data not shown). These results are expected, especially when considering that the minor differences between our assumed biomass composition and the one used in the GEM involve prototrophic amino acids (Ala, Gln, Gly) (Supplementary Figure 4).

Although the deviations are considerable in some cases, our predictions are similar to those reported for the CHO GeM (Hefzi et al., 2016), and are better for Carinhas HP, Selvarasu Late Exp, Martinez Cold 1, and Martinez Warm 2. Overall, the GeM has an average percent deviation across the ten experimental *μ*_*g,mAb*_ values of 30%, whereas our model has an average deviation of 32.5%. When considering that our model includes only 144 reactions when compared to 6,663 in the GeM, a 2.5% reduction in predictive capability is acceptable, especially considering gains in model output consistency and interpretability (to be discussed in subsequent sections) as well as reductions in computational expense, when using non-linear, multi-objective optimisation strategies for solution.

#### 3.1.1. CHOmpact identifies the source of *μ*_*g*_ prediction inaccuracies

An advantage of our proposed multi-objective optimisation strategy is that it enables the identification of nutrient uptake rates that lead to shortfalls in calculated specific growth rates when compared with the experimental values. This is achieved by constraining the maximum allowable value for the percent error of biomass growth rate and specific productivity to low values (5×10^−6^) while fixing the uptake/secretion flux values of Glc, Lac, NH_4_^+^, Pyr, Ala, Asn, Asp, Glu, Gly to the experimentally determined flux values. The upper bounds for the uptake fluxes of the remaining (mainly auxotrophic) amino acids are relaxed and obtained via constrained optimisation where the SSE is minimised. This strategy ensures that the optimal solution matches the experimental growth rate and specific productivity while finding the combination of amino acid uptake fluxes that minimises deviations from the experimental uptake rates.

The above strategy yields the results presented in Table 2, where the percent increases in specific uptake rates required for matching the experimentally determined *μ*_*g*_ and *q*_*p*_ are shown (positive/red values denote percent increase in uptake fluxes required to match *μ*_*g*_ and *q*_*p*_). Table 2 shows that for Car LP (NaBu), Selv (Early), Selv (Late), and Mart (Cold2), small or no increases in amino acid uptake rates are required to match *μ*_*g*_ and *q*_*p*_. This is expected, considering the results of Figure 2, where the predicted values for *μ*_*g*_ are matched or exceeded for these datasets.

**Table 2.**
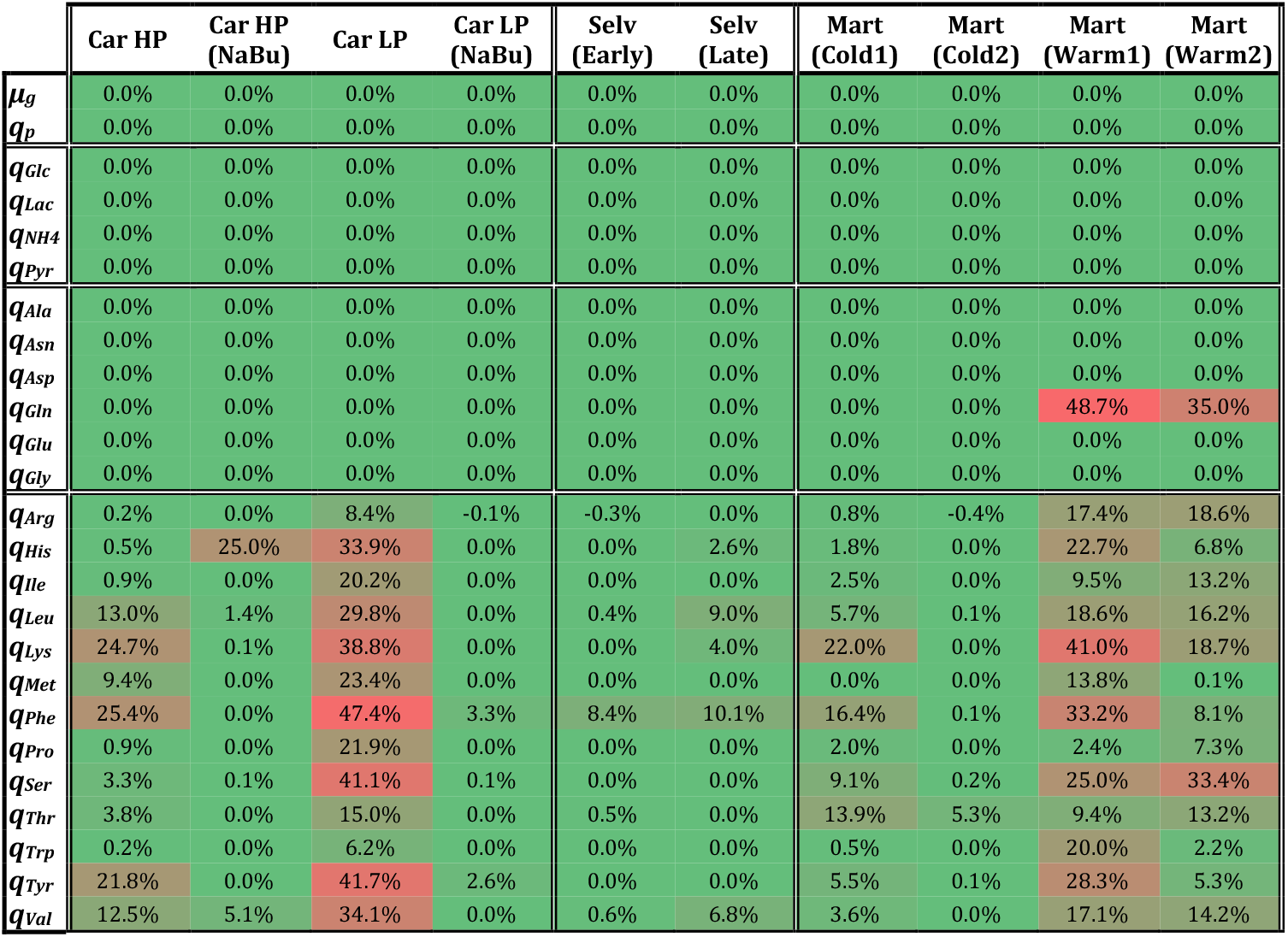
Percent increases in amino acid uptake rates required to match *μ*_g_ and *q*_*p*_. The intensity of the colours (from green = 0% to red) corresponds to magnitudes across all datasets.

|  | Car HP | Car HP (NaBu) | Car LP | Car LP (NaBu) | Selv (Early) | Selv (Late) | Mart (Cold1) | Mart (Cold2) | Mart (Warm1) | Mart (Warm2) |
| --- | --- | --- | --- | --- | --- | --- | --- | --- | --- | --- |
| $\mu_g$ | 0.0% | 0.0% | 0.0% | 0.0% | 0.0% | 0.0% | 0.0% | 0.0% | 0.0% | 0.0% |
| $q_p$ | 0.0% | 0.0% | 0.0% | 0.0% | 0.0% | 0.0% | 0.0% | 0.0% | 0.0% | 0.0% |
| $q_{Glc}$ | 0.0% | 0.0% | 0.0% | 0.0% | 0.0% | 0.0% | 0.0% | 0.0% | 0.0% | 0.0% |
| $q_{Lac}$ | 0.0% | 0.0% | 0.0% | 0.0% | 0.0% | 0.0% | 0.0% | 0.0% | 0.0% | 0.0% |
| $q_{NH4}$ | 0.0% | 0.0% | 0.0% | 0.0% | 0.0% | 0.0% | 0.0% | 0.0% | 0.0% | 0.0% |
| $q_{Pyr}$ | 0.0% | 0.0% | 0.0% | 0.0% | 0.0% | 0.0% | 0.0% | 0.0% | 0.0% | 0.0% |
| $q_{Ala}$ | 0.0% | 0.0% | 0.0% | 0.0% | 0.0% | 0.0% | 0.0% | 0.0% | 0.0% | 0.0% |
| $q_{Asn}$ | 0.0% | 0.0% | 0.0% | 0.0% | 0.0% | 0.0% | 0.0% | 0.0% | 0.0% | 0.0% |
| $q_{Asp}$ | 0.0% | 0.0% | 0.0% | 0.0% | 0.0% | 0.0% | 0.0% | 0.0% | 0.0% | 0.0% |
| $q_{Gln}$ | 0.0% | 0.0% | 0.0% | 0.0% | 0.0% | 0.0% | 0.0% | 0.0% | 48.7% | 35.0% |
| $q_{Glu}$ | 0.0% | 0.0% | 0.0% | 0.0% | 0.0% | 0.0% | 0.0% | 0.0% | 0.0% | 0.0% |
| $q_{Gly}$ | 0.0% | 0.0% | 0.0% | 0.0% | 0.0% | 0.0% | 0.0% | 0.0% | 0.0% | 0.0% |
| $q_{Arg}$ | 0.2% | 0.0% | 8.4% | -0.1% | -0.3% | 0.0% | 0.8% | -0.4% | 17.4% | 18.6% |
| $q_{His}$ | 0.5% | 25.0% | 33.9% | 0.0% | 0.0% | 2.6% | 1.8% | 0.0% | 22.7% | 6.8% |
| $q_{Ile}$ | 0.9% | 0.0% | 20.2% | 0.0% | 0.0% | 0.0% | 2.5% | 0.0% | 9.5% | 13.2% |
| $q_{Leu}$ | 13.0% | 1.4% | 29.8% | 0.0% | 0.4% | 9.0% | 5.7% | 0.1% | 18.6% | 16.2% |
| $q_{Lys}$ | 24.7% | 0.1% | 38.8% | 0.0% | 0.0% | 4.0% | 22.0% | 0.0% | 41.0% | 18.7% |
| $q_{Met}$ | 9.4% | 0.0% | 23.4% | 0.0% | 0.0% | 0.0% | 0.0% | 0.0% | 13.8% | 0.1% |
| $q_{Phe}$ | 25.4% | 0.0% | 47.4% | 3.3% | 8.4% | 10.1% | 16.4% | 0.1% | 33.2% | 8.1% |
| $q_{Pro}$ | 0.9% | 0.0% | 21.9% | 0.0% | 0.0% | 0.0% | 2.0% | 0.0% | 2.4% | 7.3% |
| $q_{Ser}$ | 3.3% | 0.1% | 41.1% | 0.1% | 0.0% | 0.0% | 9.1% | 0.2% | 25.0% | 33.4% |
| $q_{Thr}$ | 3.8% | 0.0% | 15.0% | 0.0% | 0.5% | 0.0% | 13.9% | 5.3% | 9.4% | 13.2% |
| $q_{Trp}$ | 0.2% | 0.0% | 6.2% | 0.0% | 0.0% | 0.0% | 0.5% | 0.0% | 20.0% | 2.2% |
| $q_{Tyr}$ | 21.8% | 0.0% | 41.7% | 2.6% | 0.0% | 0.0% | 5.5% | 0.1% | 28.3% | 5.3% |
| $q_{Val}$ | 12.5% | 5.1% | 34.1% | 0.0% | 0.6% | 6.8% | 3.6% | 0.0% | 17.1% | 14.2% |

Across all other datasets, substantial increases in amino acid uptake rates are required to match the experimental values for *μ*_*g*_ and *q*_*p*_. Except for both Martínez et al. (2015) Warm datasets, the uptake rate for amino acids which limit growth are auxotrophic (outlined in the bottom half of Table 2) and, in all but one case (Car HP NaBu), multiple auxotrophic amino acids limit the growth rate predictions. Particularly sharp deviations across multiple amino acids are observed for the Car LP and both Mart Warm datasets. In addition, both Mart Warm datasets are the only ones that require increased uptake of Gln.

Interestingly, when the upper bound for the Gln uptake flux is completely relaxed for the Mart Warm 2 dataset, a 74.5% increase in uptake for this nutrient is obtained, and only Lys (required 14.2% increase) and Thr (required 6.7% increase) are observed to limit growth (data not shown). This result indicates that, for this dataset, the uptake rates of the remaining amino acids do not stoichiometrically limit biomass synthesis. Rather, the limitation in growth is associated with the relatively low uptake rate of Ser which, according to our model, can be overcome by Glu-associated biosynthesis via *F*_55_. It is this additional intracellular Glu demand which pushes the uptake rates for Gln, Ile, Leu, Lys, and Val to higher values (Glu can be produced from these amino acids via reactions *F*_32_, *F*_44_, *F*_45_, *F*_39_, and *F*_41_, respectively).

Full relaxation of the Gln uptake upper bound leads to a 65.9% increase and completely curbs the excess requirements for Ser. However, the excess demand for all remaining auxotrophic amino acids (His, Ile, Leu, Lys, Met, Phe, Trp, Tyr, and Val) is unchanged, indicating that for the Mart Cold 1 dataset, the uptake rates of auxotrophic amino acids stoichiometrically limit growth.

It is key to mention that underestimations for *μ*_*g*_ arise from two possible sources (or a combination thereof): (*i*) either the assumed stoichiometric coefficients for auxotrophic amino acids in biomass are too high or (*ii*) the measured uptake rates for auxotrophic amino acids is underestimated. Beyond typical experimental variability, the measured uptake rates are unlikely to have such drastic effects on the predicted growth (errors in uptake rate measurements above 30% are unlikely – see Table 2). Therefore, uncertainty associated with biomass composition is the more likely culprit, especially when considering that it has seldom been measured for flux balance studies.

Questions surrounding CHO cell biomass composition have been recently addressed, where an average protein content (w/w) of 55.7% ± 5.5% and a dry cell weight (DCW) of 262.1 ± 28.2 pg/cell is reported for multiple CHO cell lines cultured under different conditions (Szeliova et al., 2020). When comparing these values with the 350pg/cell and 74.2% w/w protein assumed (not measured) for the datasets with the largest shortfalls in predicted *μ*_*g*_ (Martinez Warm), a maximum reduction of 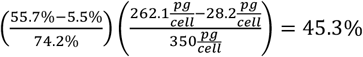 in amino acid demand towards biomass synthesis can be computed. Such a reduction would compensate for the calculated shortfalls in auxotrophic amino acid uptake rates presented in Table 2 and, therefore, enhance the predictive capability of both our model and Hefzi’s GeM.

Although detailed data for CHO cell biomass composition is now available (Szeliova et al., 2020), the values of 350pg/cell and 74.2% w/w protein were used for the results presented in Figure 2 and Table 2 in order to pinpoint how model predictive capability is impacted by our reduced reaction network when compared with the full GeM. In addition, Supplementary Figure 4 provides a comparison between the stoichiometric coefficients of amino acids in biomass assumed here with the experimentally determined ones from Szeliova et al. (2020). Despite considerable differences in DCW (219 pg/cell for exponential growth vs. 262.1 ± 28.2 pg/cell) and protein content (74.2% vs. 55.7% ± 5.5%), the stoichiometric coefficients result in quite comparable values. Therefore, we have elected to retain our originally assumed biomass composition for all subsequent calculations presented herein.

The above results highlight the importance of using accurate measurements for DCW and biomass composition in flux balance modelling. However, it is also important to mention that that biomass composition will impact model predictive capabilities only when auxotrophic components of biomass and product stoichiometrically limit growth and will be less of a factor when nutrient uptake rates exceed stoichiometric requirements. The latter case is observed for the Martinez Cold 2 dataset, where both the GeM and our model considerably over-predict *μ*_*g*_.

Several strategies have been developed to address situations where cell growth is not limited by stoichiometry. On one hand, exploring different metabolic objectives, such as maximising energetic efficiency, minimising redox stress or the uptake of essential nutrients (Chen et al., 2019; Feist and Palsson, 2010; Schuetz et al., 2007) can improve the predictive capability of flux models where nutrient uptake rates stoichiometrically exceed demand for growth and product synthesis. On another hand, strategies to further constrain the allowable values of fluxes through the reaction network have also been explored. For example, Lularevic et al. (2019) provide additional constraints using carbon balancing and Yeo et al. (2020) have developed an interesting framework whereby the fluxes through the reaction network are constrained based on the maximum rates and expression levels of the corresponding metabolic enzymes.

### 3.2. CHOmpact facilitates easier interpretation of flux distributions

Our reduced reaction network and multi-objective optimisation strategy has two key advantages over full genome-scale models and the traditional growth rate maximisation objective function used to solve them. Firstly, our multi-objective optimisation strategy enables us to calculate flux distributions across different growth phases (beyond exponential growth) and, therefore, provides enhanced insight into flux distribution dynamics. In addition, the use of multiple objectives further constrains the solution space to achieve flux distributions that are biologically consistent. Secondly, our reduced reaction network simplifies model output interpretation and allows us to better relate the obtained flux distributions with cellular physiology.

#### 3.2.1. Flux distribution dynamics

Figure 3 presents central carbon metabolism and Asp-Mal shuttle fluxes obtained with our reduced reaction network and multi-objective optimisation framework, where the flux maps correspond to three different feed compositions (Feed C, Feed U, and Feed U40) across the five phases of culture identified Supplementary Figure 2.

**Figure 3.**
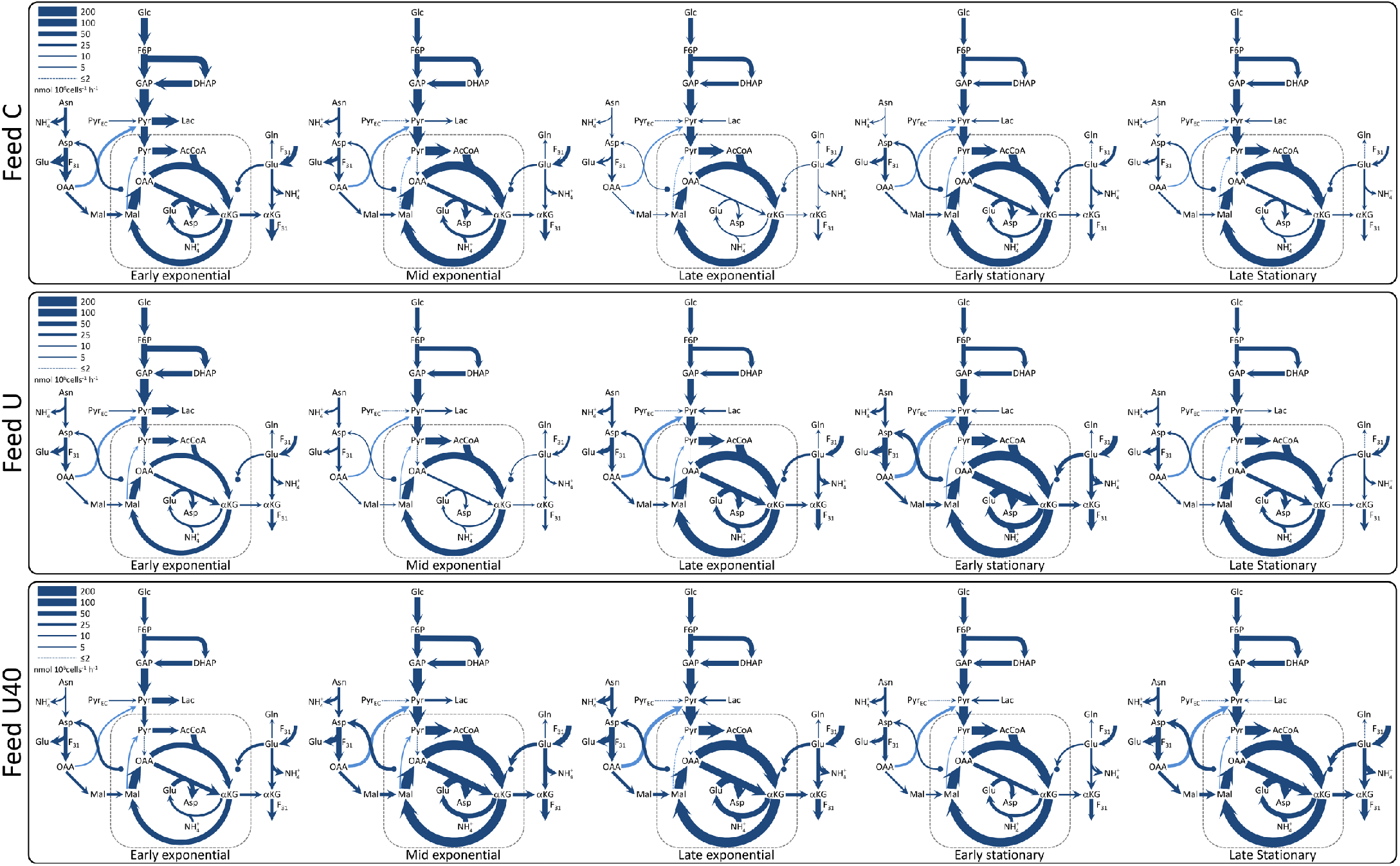
Flux map comparison. The flux distributions for central carbon metabolism and the aspartate/malate shuttle are shown for five culture intervals (early, mid, and late exponential as well as early and late stationary) across three feed compositions (Feed C, U, and U40). The line thickness corresponds to flux magnitude, as indicated by the legend at the top left-hand corner of each box.

For all feed compositions, the glycolytic fluxes decrease with time, with the highest fluxes observed during early exponential growth and the lowest during stationary growth. The magnitude of the glycolytic flux is largely determined by the glucose uptake rates, which are shown to steadily decrease as culture progresses (Supplementary Figure 2). Conversely, the tricarboxylic acid (TCA) pathway fluxes increase with culture time, which occurs because reduced lactate production allows for more glycolysis-derived pyruvate to reach mitochondria. This phenomenon, often referred to as the Warburg effect, has been widely reported for rapidly proliferating cells, including CHO cells (Buchsteiner et al., 2018; Kelly et al., 2018).

The calculated fluxes through the Asp-Mal shuttle are defined by our imposed constraint on F_20_ (Table 1), which limits its value to the flux of ‘free’ Glu (i.e., the sum of fluxes where this amino acid is produced which are not involved in the Asp-Mal shuttle). This constraint was set because all fluxes along the Asp-Mal shuttle cancel out and, therefore, can have any magnitude if the transfer of reducing equivalents from cytosolic to mitochondrial NADH is balanced. If left unconstrained, Asp-Mal shuttle fluxes have been reported to reach values that are comparable to those through central carbon metabolism (Mulukutla et al., 2012; Nolan and Lee, 2011). Although mathematically correct, these excessive flux values would be limited by the intracellular availability of Glu, which is the key substrate for the rate-limiting Asp-Mal shuttle flux (*F*_20_) (LaNoue et al., 1974; LaNoue and Tischler, 1974). Despite not directly representing Glu availability, our proposed constraint does limit the upper values of Asp-Mal shuttle fluxes to ones that fall well below those of glycolysis and, therefore, make them more biologically consistent.

#### 3.2.2. Glutamate anaplerosis and cataplerosis

Glutamate can either be consumed towards TCA and energy production (anaplerosis) or produced from TCA metabolites for subsequent use in biomass generation (cataplerosis). According to our reaction network, net Glu anaplerosis occurs when more of this amino acid is transported into mitochondria (*F*_20_) than what is transported out, in the form of αKG. Net Glu cataplerosis occurs when less of it is transported into mitochondria than the αKG transported out. αKG is then converted into Glu by either *F*_30_ or *F*_31_, the latter of which uses Asp as a co-substrate.

Figure 4 shows that, in Feed C, net Glu cataplerosis (Glu produced from TCA) is observed during the three exponential growth intervals. This trend is reversed during stationary phase, where net anaplerosis (Glu consumed towards TCA) occurs. Feed U and Feed U40 contrast with Feed C in that they present Glu anaplerosis (consumption towards TCA) across all but one culture interval (Figure 4 – bottom).

**Figure 4.**
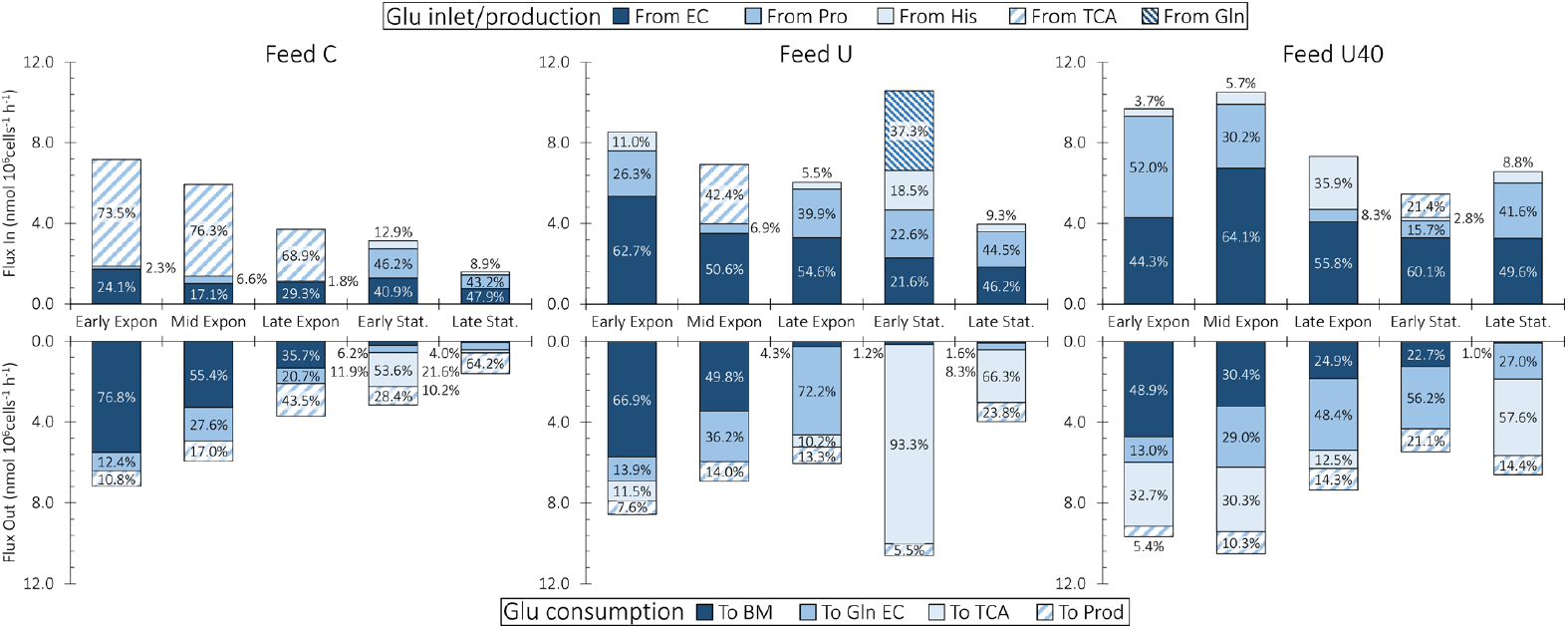
Glutamate flux distributions. The glutamate inlets for all feed compositions are shown in the top half and the outlets are shown in the bottom half. Glutamate sources and sinks are indicated by the shading and the top and bottom legends. The percent contributions to the total inlet or outlet are shown within the bars.

In Feed U, Glu cataplerosis is only observed during mid exponential growth, where its production from TCA accounts for 42.4% of the total Glu inlet (Figure 4 – top). Another interesting feature from Feed U is the level of anaplerosis observed during the early stationary interval, where it accounts for 93.3% of the total consumed Glu (Figure 4 – bottom). This high level of anaplerosis is associated with increased Glu production from Gln (37.3% of total) and, to a lesser extent, from His (18.5% of total) (Figure 4 – top). It is worth noting that this is the only culture condition and interval where Gln is consumed from the extracellular environment (Supplementary Figure 1).

In Feed U40, Glu cataplerosis is observed only during the early stationary interval, where 21.4% of all Glu produced is derived from TCA (Figure 4 – top). Glu cataplerosis during early stationary phase is most likely to arise to offset the high level of Gln secretion observed during this interval (Supplementary Figure 1).

In our GS-CHO Feed C cultures, Glu cataplerosis is driven by the high Asn uptake rates observed during exponential growth (Supplementary Figure 2), where a high *F*_31_ flux provides a path for Asn overflow towards pyruvate via oaxaloacetate (*F*_29_). An alternative cataplerotic pathway for Glu is its direct synthesis from cytosolic αKG via *F*_30_ (catalysed by enzyme <u>EC 1.4.1.3</u>), which consumes NH_4_^+^ and produces cytosolic NAD^+^ (Yang et al., 2005) and may, thereby, reduce lactate production (Freund and Croughan, 2018). Its ability to reduce NH_4_^+^ and lactate production make <u>EC 1.4.1.3</u> a potential target for metabolic engineering.

Prior flux balance work on standard (non-GS) CHO cells commonly reports net Glu anaplerosis during the exponential growth phase of cells, where high lactate production results in low TCA fluxes (Ahn and Antoniewicz, 2013). It is thought that Glu anaplerosis is used by the cells to replenish flux through TCA and is also known to be a major source of ammonia production because much of the anaplerotic flux involves glutamine aminolysis (Dean and Reddy, 2013; Wahrheit et al., 2014).

In contrast to past work, our FBA results indicate substantial Glu anaplerosis during the stationary phases of culture across all three feeding strategies (Figure 4 – bottom). Our results suggest that the high uptake rate of Asn observed across all cultures throttles Glu anaplerosis. The Asn overflow pathway discussed above produces Glu in F_31_, which, in turn causes Glu overflow that is taken up by TCA. Anaplerosis as a means to cope with Glu overflow is also substantiated by the early stationary phase flux distribution of Feed U, which presents the highest level of Glu anaplerosis observed across all datasets (9.88 nmol/10^6^cells/h) (Figure 4 – bottom). This culture interval is the only one across all datasets where glutamine (Gln) is consumed by the cells. It is this Gln uptake which causes Glu overflow through the glutaminolysis pathway that is commonly reported for non-GS-CHO cells.

Glu anaplerosis is commonly described as the cytosolic production of αKG from Glu (via *F*_30_ – Figure 1), the former of which is transported into the mitochondrial matrix (*F*_21_) for uptake by TCA (Ahn and Antoniewicz, 2013; Mulukutla et al., 2012; Nicolae et al., 2014). Crucially, the transport of αKG into the mitochondrial matrix depends on malate availability in the cytosol: *F*_21_ (OGCP) has an antiport mechanism whereby one molecule of αKG is transported into the mitochondrial lumen for every malate molecule that is transported out (Iacobazzi et al., 1992).

Our results indicate an alternative mechanism for Glu anaplerosis, where Glu is first transported into the mitochondrial matrix (via *F*_20_ – the rate-limiting and only irreversible reaction of the Asp-Mal shuttle (LaNoue et al., 1974; LaNoue and Tischler, 1974)) where it is then converted, with the consumption of mitochondrial Asp, into αKG via *F*_22_. This alternative mechanism arises from the high Asn uptake by our GS-CHO cells, where a substantial amount of this nutrient is funnelled towards cytosolic malate through reactions *F*_51_, *F*_31_, and *F*_24_. This Asn overflow pushes αKG out of the mitochondrial matrix where it can be consumed to produce Glu, mainly through *F*_31_. Irrespective of the metabolic route, the magnitudes of Glu anaplerosis and cataplerosis are consistent with those determined through Metabolic Flux Analysis (Ahn and Antoniewicz, 2013; Nicolae et al., 2014).

#### 3.2.3. Asparagine and aspartate are key anaplerotic nutrients

The asparagine/aspartate pair (Asn/Asp), linked through *F*_51_, is a key contributor to TCA flux through its sequential conversion to oxaloacetate (*F*_31_) and pyruvate (*F*_29_), the latter of which is ultimately transported into mitochondria for consumption in the TCA. Because alanine and lactate are also produced from pyruvate (through reactions *F*_26_ and *F*_25_, respectively), the Asn/Asp pair is also a key source of these metabolites.

During the early exponential interval, cells cultured with Feed C consume 43.9% of Asn/Asp towards TCA and 34.1% towards lactate (Figure 5 – bottom). As lactate secretion subsides, the contribution of Asn/Asp towards TCA increases to beyond 70% for the remaining culture intervals. The trend for Asn/Asp consumption towards TCA is even more pronounced for cells cultured with Feeds U and U40, where it exceeds 85% across the final three culture intervals.

**Figure 5.**
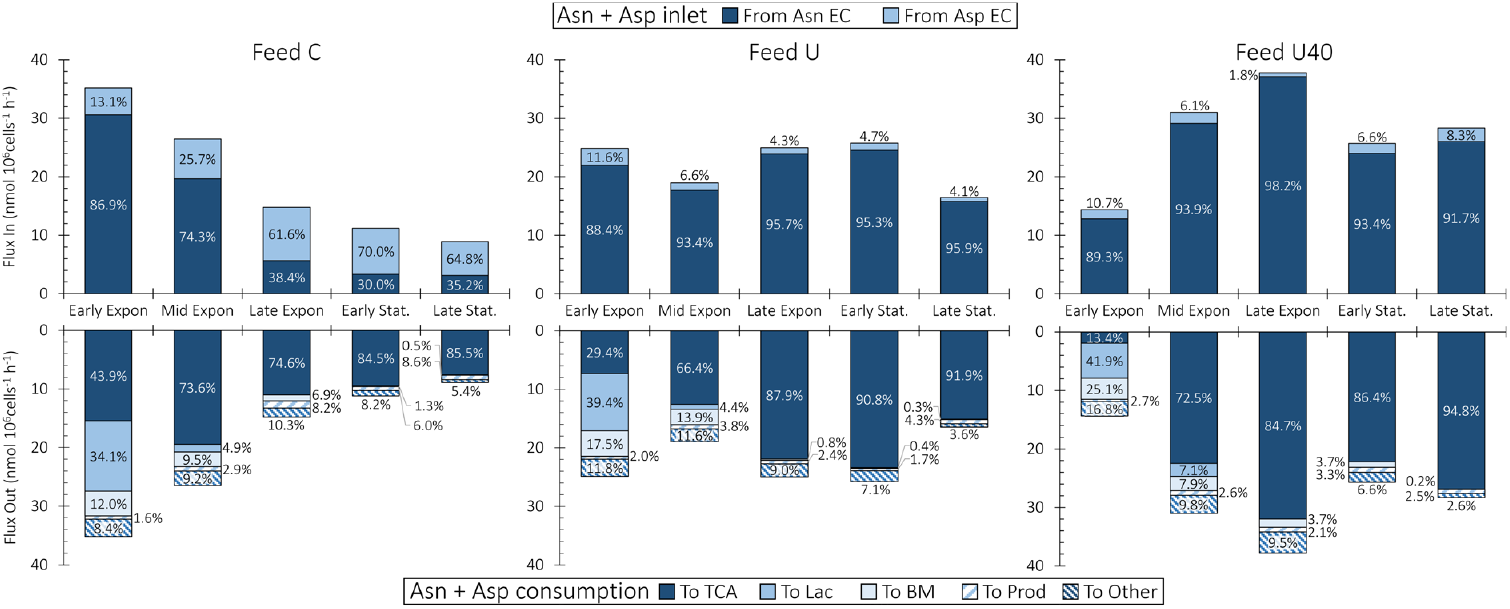
Asparagine + Aspartate flux distributions. The sources for asparagine (Asn) and aspartate (Asp) for all feed compositions is shown in the top half and the sinks are shown in the bottom half. The source and sink fluxes for Asn and Asp have been summed for simplicity. The different Asn+Asp sources and sinks are indicated by the shading and the top and bottom legends. The percent contributions to the total sources and sinks are shown within the bars.

Across all feeds, the maximum proportion of Asn/Asp consumed towards biomass and mAb product is 27.8% for Feed U40 during early exponential phase (Figure 5 – bottom). These results show that Asn/Asp are fed well beyond stoichiometric requirements for growth and product formation and that anaplerosis provides an overflow pathway for when these nutrients are fed in excess. These results are consistent with prior work where excess Asn/Asp feeding has been observed to increase alanine and lactate secretion by CHO cells (Calmels et al., 2019; Selvarasu et al., 2012).

The anaplerotic overflow pathway for Asn/Asp results in a rapid uptake of Asn and a concomitant reduction in its concentration in the culture medium (Supplementary Figure 1). Particularly low residual Asn concentrations are observed in the Feed C culture, where Asn and Asp are fed at the lowest levels. In absence of flux balance calculations, the observed reduction in Asn availability could be interpreted as being growth limiting and would lead to increasing the concentration of Asn in the media and/or feed to alleviate the perceived bottleneck. Our experimental and FBA results demonstrate that Asn is indeed not a growth limiting nutrient, and that increasing its concentration in the feed in fact reduces cell growth (Supplementary Figure 1), likely due to the production of ammonia associated with Asn anaplerosis. Similar observations were recently reported by Calmels et al. (2019), where GeM FBA calculations were applied to industrial CHO DG44 cells.

FBA also allows us to estimate the Asn/Asp uptake rates at which the anaplerotic pathway becomes saturated (i.e., where no more Asn/Asp can be funnelled towards TCA). The raw experimental data shows that Asp accumulates in the extracellular environment of Feed U40 cultures during the late exponential, early stationary, and stationary phases (Supplementary Figure 1) implying that, under these conditions, the cells resort to secreting Asp instead of consuming it towards central carbon metabolism. The flux of Asn/Asp towards TCA during these culture phases are 38.7 nmol/10^6^cells/h, 25.7 nmol/10^6^cells/h, and 32.1 nmol/10^6^cells/h, respectively and represent the range of Asn/Asp overflow our GS-CHO cells can cope with. Interestingly, these values closely correlate with the values of total available Glu flux, which are 7.3 nmol/10^6^cells/h, 5.5 nmol/10^6^cells/h, and 6.6 nmol/10^6^cells/h for the corresponding culture intervals. This correlation indicates that intracellular Glu availability may regulate the extent of Asn/Asp anaplerosis.

#### 3.2.4. NH_4+_ sources and sinks

NH_4_^+^ is a key determinant of CHO cell culture performance because it is known to impact cell growth (Synoground et al., 2021; Wahrheit et al., 2014) as well as product quality (Borys et al., 1994; Hong et al., 2010). In standard CHO cells, NH_4_^+^ is mainly generated as a by-product of Gln anaplerosis (glutaminolysis) (Dean and Reddy, 2013; Hong et al., 2010; Wahrheit et al., 2014). Glutamine synthase (GS) cells, such as the ones used in this study, satisfy their Gln requirements by producing it from Glu via ectopic expression of glutamine synthase, thus circumventing the negative impact of NH_4_^+^ on the cell culture process. Despite considerable reductions in NH_4_^+^ accumulation, GS-CHO cells still produce ammonia to levels that may still impact product glycosylation (Borys et al., 1994; Hong et al., 2010) so it is therefore important to characterise the major sources and sinks of this key metabolite.

The dark blue bars in the top half of Figure 6 show that the vast majority (>75%) of ammonia is produced from asparagine (*F*_51_). The only exceptions are the Early Stationary intervals of Feed C and Feed U where, respectively, Asn is the source of 46.4% and 67.5% of all produced NH_4_^+^. During the Early Stationary phase of Feed C, the lower levels of NH_4_^+^ production from Asn are due to depletion of this nutrient in the culture media along with NH_4_^+^ uptake by the cells (Supplementary Figure 1). In Feed U, the lower proportion of NH_4_^+^ generated from Asn is caused by glutaminolysis – this is the only interval across all experiments where Gln is consumed by the cells (Supplementary Figure 1). Additional sources of NH_4_^+^ include Ser (*F*_36_), Thr (*F*_37_) and His (*F*_54_), although to much lower levels, when compared with Asn. These results indicate that, in GS-CHO cells, NH_4_^+^ production is throttled by the Asn/Asp anaplerosis discussed in section 3.2.3 and is consistent with previous work with GS-CHO cells (Calmels et al., 2019; Carinhas et al., 2013).

**Figure 6.**
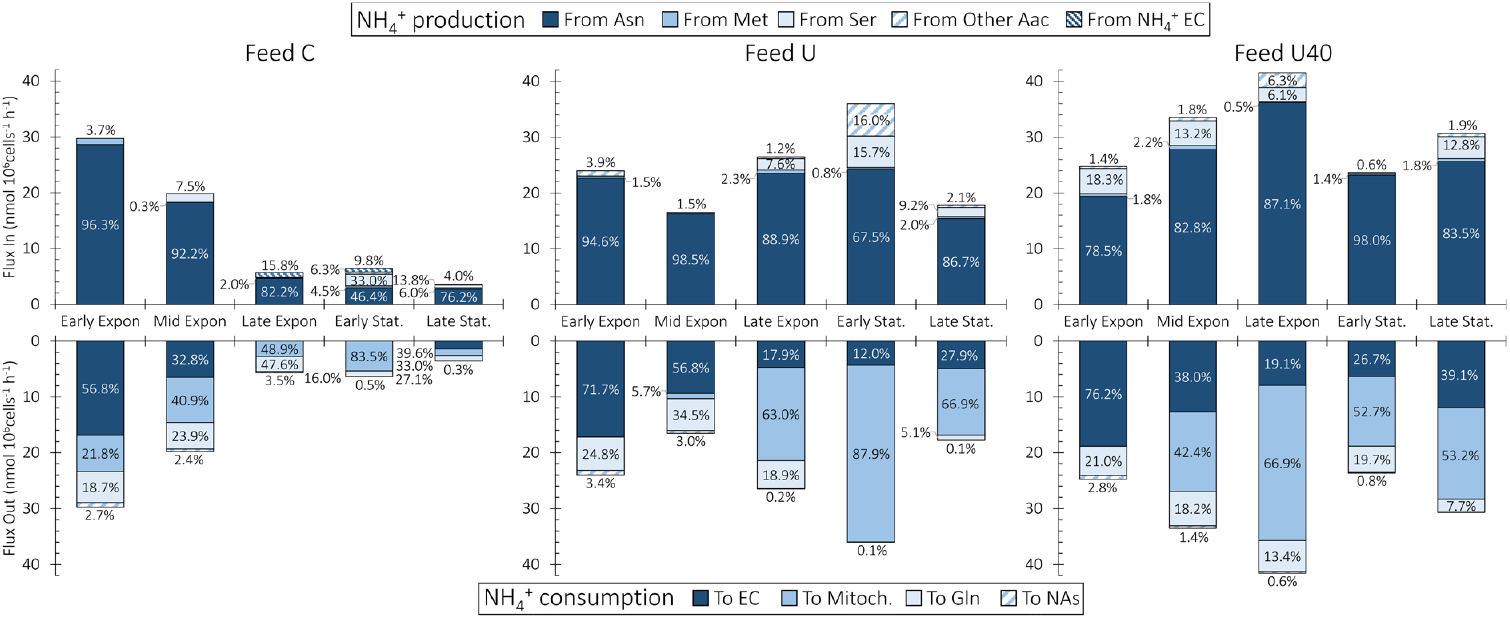
NH_4_^+^ Flux distributions. The NH_4_^+^ sources for all feed compositions are shown in the top half and the sinks are shown in the bottom half. The different sources and sinks are indicated by the shading and the top and bottom legends. The percent contribution of each source/sink to the total produced/consumed is shown within the bars.

The total production rate of NH_4_^+^ further confirms its link with Asn/Asp anaplerosis. The top half of Figure 6 shows that the total amount of NH_4_^+^ produced by the cells increases with higher levels of Asn feeding. After the Mid Exponential interval, cells cultured with basal Asn feeding levels (Feed C) have NH_4_^+^ production rates below 8 nmol/10^6^cells/h, whereas the Feed U and Feed U40 cultures (increasingly higher levels of Asn feeding) produce above 20 nmol/10^6^cells/h of NH_4_^+^.

The bottom half of Figure 6 presents the major NH_4_^+^ sinks across the three feed conditions, as obtained through our FBA framework. There, it can be seen that three NH_4_^+^ sinks predominate: (*i*) secretion to the extracellular (EC) environment (dark blue bars – F_108_), (*ii*) consumption in mitochondria (intermediate blue bars) and (*iii*) consumption towards Gln synthesis (light blue bars – F_32_). Of the three major sinks, EC secretion can be deduced from the experimental data. The sink towards Gln synthesis is similarly intuitive and occurs because our GS-CHO cells are not fed Gln and must cover their demand for this amino acid through its GS-enabled synthesis using Glu and NH_4_^+^ as substrates.

The less intuitive NH_4_^+^ sink is the one associated with mitochondria. According to our FBA results, a substantial amount of NH_4_^+^ is consumed by a mitochondrial reaction cycle, where OAA is combined with Glu to produce αKG and Asp (*F*_22_) and where the resulting αKG is combined with NH_4_^+^ to produce Glu (*F*_23_). This mitochondrial NH_4_^+^ sink is governed by the presence of Asp within the mitochondrial lumen and is therefore coupled with the Asp-Mal shuttle, where *F*_20_ transports Asp out of mitochondria. If Asp accumulates within mitochondria, *F*_22_ and *F*_23_ will be reversed and may, thereby, cause net NH_4_^+^ production by mitochondria. These results are consistent with experimental findings where high NH_4_^+^ concentrations increased the cell specific consumption rates of Asp and Glu (Lao and Toth, 1997).

Due to the constraint imposed on *F*_20_ by our FBA solution strategy (discussed in Section 2.2.2 and presented in Table 1), a second sink for mitochondrial Asp is required to drive mitochondrial NH_4_^+^ consumption. Within our FBA reaction network, this additional sink is given by *F*_59_ of the urea cycle, where mitochondrial Asp is irreversibly combined with citrulline to produce fumarate and arginine (Supplementary Table 1). Our results indicate that *F*_59_ consumes over half of the mitochondrial Asp across all culture conditions (Supplementary Figure 5) and is, therefore, a key determinant of mitochondrial consumption of NH_4_^+^.

The above mitochondrial Asp sink (*F*_59_) enables mitochondrial consumption of NH_4_^+^ that is independent of the Asp-Mal shuttle and, therefore, of cytosolic Glu availability. This Glu-independent NH_4_^+^ detoxification pathway would require diverting aKG directly from TCA to be consumed in reaction *F*_23_. The produced Glu would then react with TCA-derived oxaloacetate to replenish aKG and produce mitochondrial Asp (*F*_22_). Finally, mitochondrial Asp would be consumed by *F*_59_ to yield arginine and fumarate, which could also feed back into TCA.

Interestingly, CHO cells are reported to produce only trace amounts of urea (Zamorano et al., 2010), indicating that certain enzymes of the cycle may be inactive. The mitochondrial sink identified by our FBA may be an alternate route of NH_4_^+^ detoxification (independent of urea secretion) that leverages two of the urea cycle enzymes (<u>EC 6.3.4.5</u> and <u>EC 4.3.2.1</u>) which are known to be expressed in CHO cells (Heffner et al., 2020).

## 4. Concluding remarks

We have presented a reduced reaction network to describe the metabolism of mAb-producing CHO cells. Our reduced metabolic network (144 reactions) performs comparably with the iCHO1766 GeM (>6000 reactions) in predicting the growth rates of different CHO cell lines. Our FBA framework also allowed us to identify the absence of cellular weight and composition measurements as the most likely cause of inaccuracies in predicting the growth rates.

We have also presented a comprehensive multi-objective optimisation strategy to solve our metabolic model. Our multi-objective optimisation framework constrains the solution space to yield physiologically consistent flux distributions across all phases of cell culture.

When coupled with multi-objective optimisation, our compact reaction network greatly enhances the interpretability of metabolic flux distributions across the different phases of cell culture. In this context, our results provide insights into the mechanisms underlying Glu anaplerosis and its dependence on the uptake rate of Asn/Asp. We have also identified Asn and Asp as the key anaplerotic nutrients of GS-CHO cells and that, in this role, they are an important source of lactate during the early stages of culture.

Our results also show that Asn is the predominant source of NH_4_^+^ across all culture conditions and that the major sink for this key metabolite is consumption within mitochondria. The presence of Asp within mitochondria determines whether this organelle is a source or sink of NH_4_^+^: when Asp accumulates, mitochondria can become a net source of NH_4_^+^, when Asp is depleted, NH_4_^+^ is consumed within mitochondria. The Asp-Mal shuttle determines the intracellular flux distributions of Asn, Asp, Gln, Glu and NH_4_^+^. Our FBA solution strategy constrains fluxes through the Asp-Mal to not exceed the flux of ‘free’ Glu entering or produced by the cells in order to obtain physiologically consistent flux distributions for Asn, Asp, Gln, Glu and NH_4_^+^.

Moving forward, the enhanced understanding of metabolic dynamics afforded by our reduced reaction network and multi-objective optimisation framework can be used to define feeding strategies that optimise cell culture performance. Furthermore, the compact size of our reaction network will also facilitate the creation of hybrid dynamic FBA/culture dynamics models which can be used as digital twins for dynamic optimisation and control of cell culture bioprocesses.

## Acknowledgements

CK and IJV gratefully acknowledge funding by BRIC/BBSRC. IJV also acknowledges funding from Science Foundation Ireland (12/RC/2275_P2). This manuscript is dedicated in memory of Aoife Carney, Administrative Officer, School of Chemical & Bioprocess Engineering, University College Dublin.

