## Supplementary File for "CHOmpact: a reduced metabolic model of Chinese hamster ovary cells with enhanced interpretability"

**Aspartate-Malate Shuttle**

The purpose of the Asp-Mal shuttle is to transfer reducing equivalents (electrons) from cytosolic NADH to mitochondrial $NAD^+$ (Borst, 2020; Zagari et al., 2013b). Because the mitochondrial membrane is impermeable to electrons, the following reactions are required for their transfer from cytosol to mitochondria:

| | |
|---|---|
| (+1) $F_{15}$ | $Mal_{mit} + NAD^{+}_{mit} \leftrightarrow Oxal_{mit} + NADH_{mit}$ |
| (+1) $F_{20}$ | $Glu_{cyt} + Asp_{mit} \rightarrow Glu_{mit} + Asp_{cyt}$ |
| (+1) $F_{21}$ | $Mal_{cyt} + \alpha KG_{mit} \leftrightarrow Mal_{mit} + \alpha KG_{cyt}$ |
| (+1) $F_{22}$ | $Oxal_{mit} + Glu_{mit} \leftrightarrow \alpha KG_{mit} + Asp_{mit}$ |
| (+1) $F_{24}$ | $Oxal_{cyt} + NADH_{cyt} \leftrightarrow Mal_{cyt} + NAD^{+}_{cyt}$ |
| (+1) $F_{31}$ | $Asp_{cyt} + \alpha KG_{cyt} \leftrightarrow Oxal_{cyt} + Glu_{cyt}$ |
| Sum | $NADH_{cyt} + NAD^{+}_{mit} \rightarrow NADH_{mit} + NAD^{+}_{cyt}$ |

From the above, it can be seen that the Asp-Mal shuttle reactions yield no net production/consumption of Asp, Glu, αKG, Oxal or Mal because, within this pathway, all participating components (except $NADH_{cyt}$, $NAD^{+}_{mit}$, $NADH_{mit}$, and $NAD^{+}_{cyt}$) are produced and consumed at equivalent rates – their fluxes within the shuttle cancel out. In conjunction, the sink for mitochondrial $NADH_{mit}$ is unconstrained: it is consumed to produce ATP via $F_{19}$ and the flux of ATP consumption towards unspecified reactions ($F_{104}$ – see Figure 1 in the manuscript) is computed through optimisation.

Overall, this means that as long as the balances for $NADH_{cyt}$, $NAD^{+}_{mit}$, $NADH_{mit}$, and $NAD^{+}_{cyt}$ are satisfied, the fluxes for all reactions in the Asp-Mal shuttle can take any value and may lead to shuttle uptakes of glutamate that are unfeasibly high (e.g., having similar magnitude as glycolytic fluxes while glutamate uptake from the extracellular environment is an order of magnitude below that of glucose).

For this reason, we have constrained $F_{20}$, the rate-limiting step of the Asp-Mal shuttle (LaNoue et al., 1974; LaNoue and Tischler, 1974), to never exceed the flux of glutamate internalised by the cells or produced through reactions independent of the Asp-Mal shuttle.

### Cell culture data

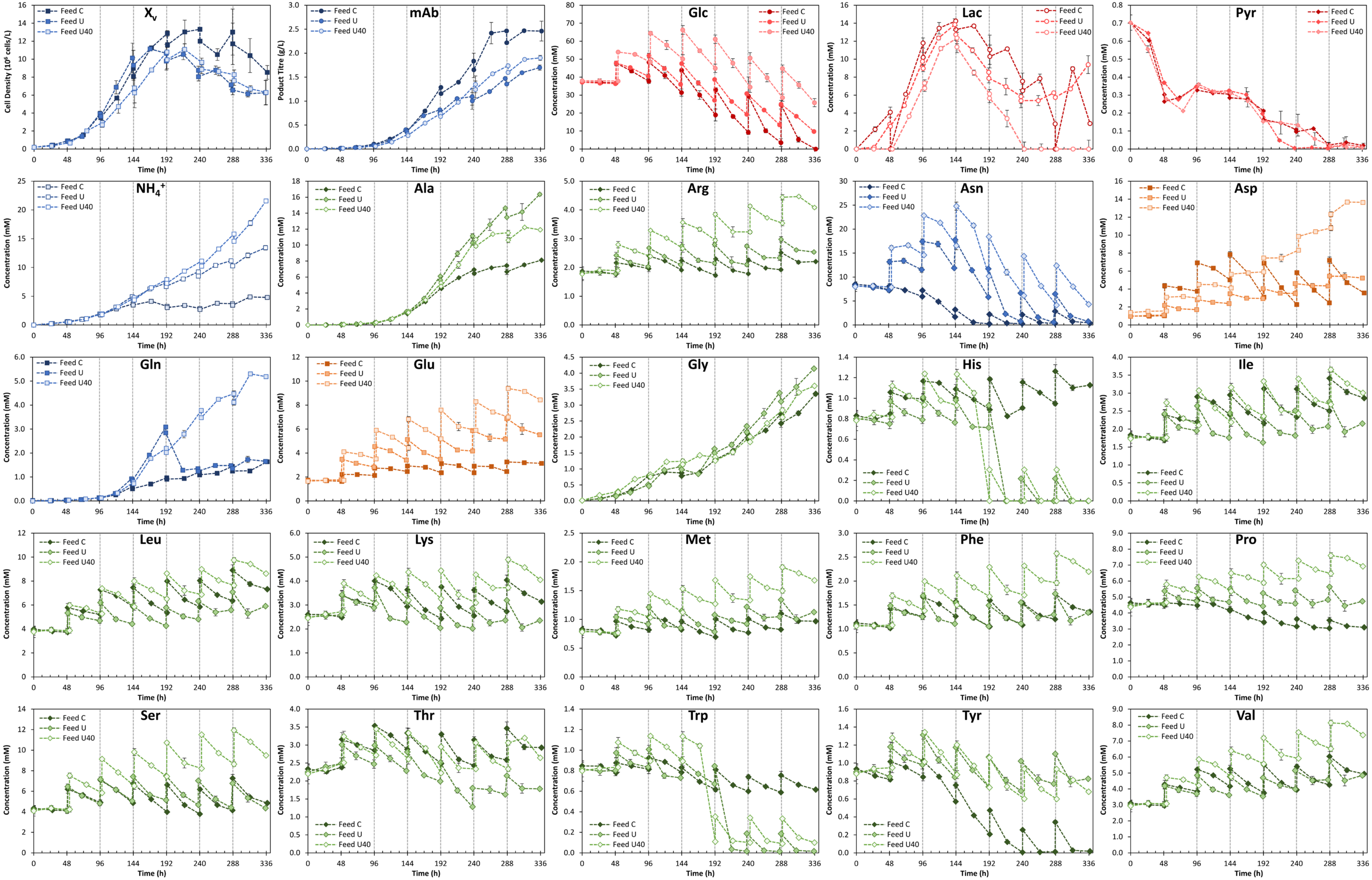


**Supplementary Figure 1. Experimental data for three fed-batch experiments. The vertical grey dashed lines show the culture intervals considered for FBA calculations.**

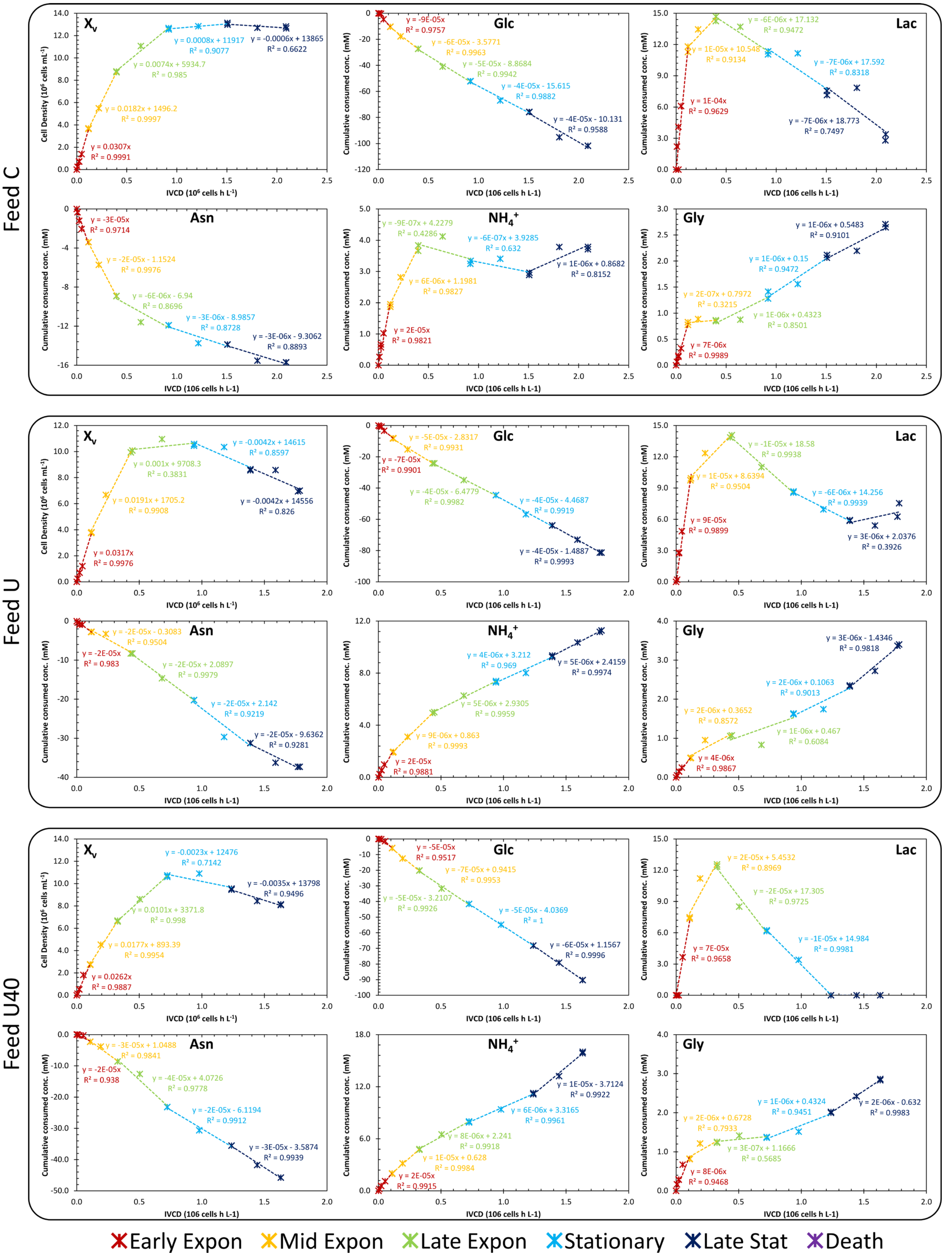


**Supplementary Figure 2. Intervals considered for flux analysis**

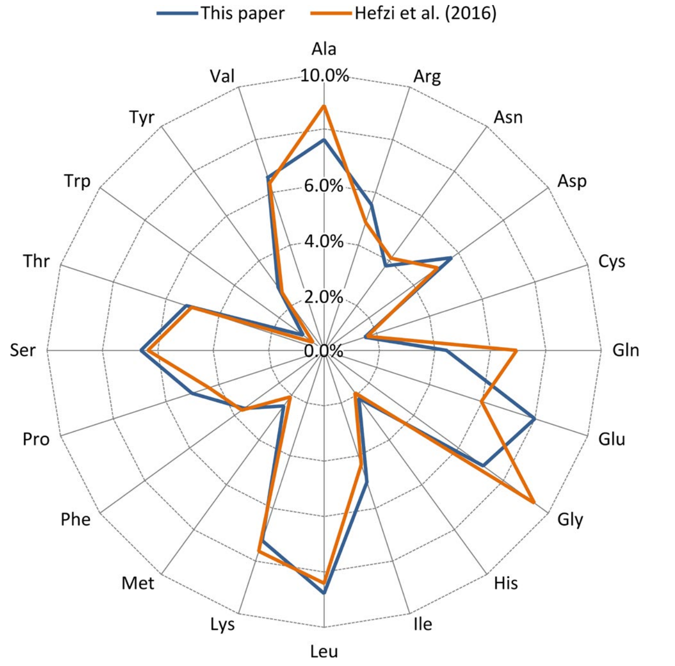


**Supplementary Figure 3. Comparison of the amino acid composition in HCPs.**
**The values considered in this paper are shown in blue and the ones assumed by Hefzi et al. (2016) are presented in orange. The composition is shown in units of mole of amino acid *i* per mole of host cell protein.**

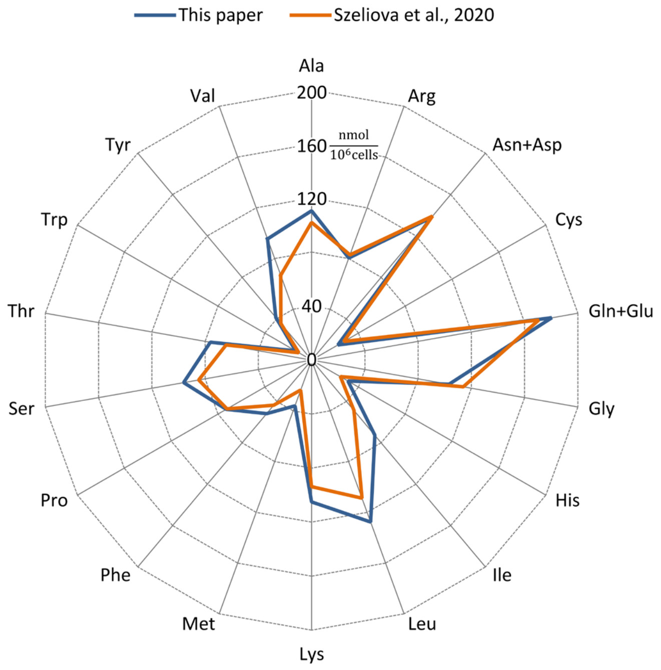


**Supplementary Figure 4. Comparison of the stoichiometric coefficients for amino acids in biomass.**
**The exponential growth stoichiometric coefficients of amino acids in biomass used in this work are compared with those determined by Szeliova et al. (2020). The latter correspond to the average value across ten CHO cell lines cultured under different nutrient conditions.**

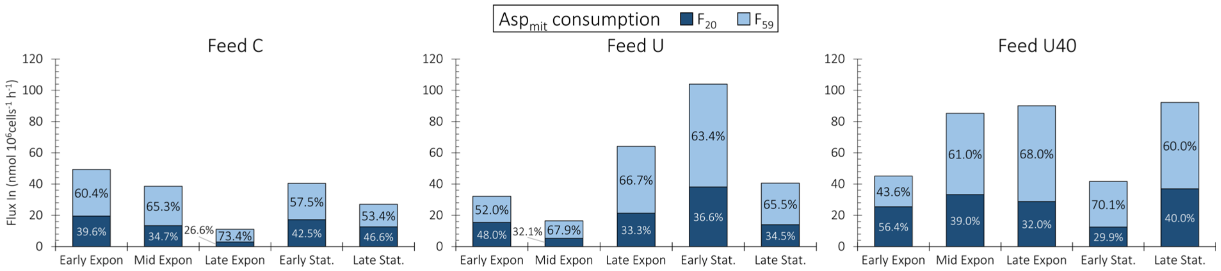


**Supplementary Figure 5. The rate-limiting step of the Asp/Mal shuttle ($F_{20}$) and the sum of cell-specific uptake rates for Asn, Asp, Gln, and Glu for the three feed compositions (Feed C, U, and U40). Similarities between the profiles for $F_{20}$ and the sum of anaplerotic amino acid uptake rates are observed. The minimum values of $F_{20}$ corresponds to a shift from Glu cataplerosis to anaplerosis.**
